# RsaM is not a switch but a built-in modulator of quorum sensing in *Pseudomonas fuscovaginae*

**DOI:** 10.64898/2026.01.05.697732

**Authors:** Nemanja Ristović, Iris Bertani, Gianluca Triolo, Michael P. Myers, Cristina Bez, Vittorio Venturi

## Abstract

*Pseudomonas fuscovaginae*, a wide host-range plant pathogen of several cereal and grass species, possesses two canonical *N*-acyl homoserine (AHL)-based quorum sensing (QS) systems called PfsI/R and PfvI/R, both of which are inactive under laboratory conditions but active *in planta*. A Tn*5* mutant insertion in the *pfsI* and *pfsR* intergenic region was previously reported to trigger the PfsI/R system. This region contains coding sequences for RsaM, a putative protein that has since been hypothesized to play a central role in imposing repression on the PfsI/R system. Putative *rsaM* genes/RsaM proteins negatively controlling AHL QS systems have also been reported in several other bacterial species. In the present study, we report for the first time the endogenous expression of an RsaM family protein and determine its position within the PfsI/R regulatory circuit. We found that RsaM is not produced in the wild-type *P. fuscovaginae* and does not play a role in keeping the PfsI/R system in a quiescent state. The expression of RsaM is instead triggered upon targeted mutations in the *pfsR*-*rsaM* intergenic region, which concomitantly activate the transcription of both the *pfsI* and *pfsR* genes. Moreover, we demonstrated that RsaM attenuates the PfsI/R circuit upon its activation. Taken together, our results evidenced that RsaM does not function as a repressor switch of the PfsI/R system, but behaves as a built-in modulator that prevents overactivation of this circuit once it is triggered.

**Importance:** *Pseudomonas fuscovaginae* is a globally occurring plant pathogen that employs AHL QS to regulate virulence. In this bacterium, QS signalling circuits display a rather unusual feature; the lack of activation at high cell densities under standard laboratory conditions. A hypothetical regulator named RsaM was previously linked to this phenomenon as a possible repressor switch acting on the PfsI/R AHL QS system in the absence of an unknown signal or stimulus. In this study, we demonstrated that the expression of RsaM and activation of the PfsI/R system both depend on the disruption of the *rsaM* and *pfsR* divergent intergenic region, and that RsaM functions as a negative modulator of this circuit, rather than as its master repressor. This study unveils the functional position of a novel protein regulator and broadens our understanding of regulatory configurations governing QS circuits.

## Introduction

*Pseudomonas fuscovaginae* is a seed-borne broad host range phytopathogen associated with brown sheath rot of rice and other cereals. It is capable of colonizing various plant niches, both as an endophyte and an epiphyte, and displays considerable plasticity in its life cycle (Rott, 1989; Bigirimana et al., 2015). Several virulence factors and virulence-associated processes, including phytotoxins, secretion systems, and quorum sensing (QS), contribute to its pathogenicity (Batoko et al., 1997; Mattiuzzo et al., 2011; Patel et al., 2014).

QS is a bacterial cell-cell communication process mediated by diffusible signalling molecules that links cell density and environmental stimuli to transcriptional regulation of gene expression, enabling coordination of various social behaviors. Among Pseudomonadota (formerly Proteobacteria), the most common QS systems consist of signalling and sensing modules, comprised of a LuxI family synthase, the producer of signalling molecules *N*-acyl homoserine lactones (AHLs), and a LuxR family transcriptional regulator, the receptor of cognate AHL(s). QS typically relies on constitutive production of AHLs, which accumulate as cell density increases. Once critical AHL concentrations are reached, these signals are detected by the LuxRs, which then dimerize and bind the *lux* boxes, the *cis*-regulatory elements located within the promoter regions of LuxR target genes. The key regulatory target of the LuxRs is most commonly the cognate *luxI* gene, setting off an autoinducing loop. This circuitry therefore results in signal amplification, gene expression synchronization across the bacterial population, and a shift from individual to group behaviors. LuxR regulons are diverse, encompassing various genes involved in the production of common goods, and thus enabling coordination of distinct collective endeavors (Miller & Bassler, 2001; Whitehead et al., 2001; Waters & Bassler, 2005). AHL-based QS systems in bacterial phytopathogens often regulate communal phenotypes involved in pathogenicity, such as motility, virulence factor production, biofilm formation, and oxidative stress tolerance (Quiñones et al., 2005; Hussain et al., 2008; Põllumaa et al., 2012).

The timing and magnitude of QS activation usually do not align only with cell density but are also affected by numerous extracellular and intracellular cues (Kindler et al., 2019; Striednig & Hilbi, 2022; Schuster et al., 2023). These signals are conveyed via protein and RNA regulators that act at the transcriptional, post-transcriptional or more rarely, at the post-translational level, and affect the amount or activity of the LuxR and LuxI proteins. While some regulators of the LuxI/R systems function globally and link QS to other regulatory cascades, others feature as intrinsic/built-in elements, typically counteracting the LuxR-driven positive feedback loop. QS is thus today regarded as a complex regulatory network that integrates a diverse array of cues beyond cell density (Venturi et al., 2011; Frederix & Downie, 2011; Spacapan et al., 2023).

*P. fuscovaginae* UPB0736 possesses two canonical AHL-based QS modules, named PfvI/R and PfsI/R. Strikingly, neither system responds to increasing cell density under standard laboratory conditions. However, the knockout mutants of the synthase or regulator genes display impaired virulence in rice and quinoa, providing evidence that AHL QS in *P. fuscovaginae* is activated during plant infection. These findings suggested involvement of other cues and regulatory pathways which, combined with AHL concentration, unlocked one or both QS systems. In a genetic screen of a Tn*5* genomic mutant library of *P. fuscovaginae* UPB0736, a mutant was identified having a transposon insertion in the intergenic region between the *pfsI* and *pfsR* genes which displayed a dramatic increase in *pfsI* promoter activity and overproduced AHLs. Two possible open reading frames (ORFs) encoding a hypothetical protein named RsaM, which are in frame with one another, were identified in this intergenic region (Fig. 1A) (Mattiuzzo et al., 2011). It was therefore hypothesized that RsaM could be a stringent repressor of the PfsI/R system that is likely to act as a switch in response to a yet unknown signal/s. The sequence of RsaM was found to be unique, with no significant homology to any characterized proteins. Similar ORFs have subsequently been identified in a number of species; interestingly the putative *rsaM* gene was always located adjacent to AHL QS loci (Venturi et al., 2023). An RsaM homologue from *Burkholderia cenocepacia*, BcRsaM, was heterologously expressed and characterized in structural and biochemical studies that evidenced this protein possesses a distinct domain architecture. Several sequence-and domain-organization features appear to be maintained across the so far predicted RsaMs, with the hallmark being the presence of a preserved tryptophane quartet localized in the protein interior (Michalska et al., 2014; Venturi et al., 2023). While all these findings suggested that *rsaM* ORFs are endogenously functional and expressed as proteins, no validation studies have been performed to date. The molecular mode of action of RsaMs, if expressed natively, could not be inferred from the studies on BcRsaM, as this protein lacks known DNA- or RNA-binding motifs, enzyme active site signatures, and is unable to bind AHLs (Michalska et al., 2014). AHL production in *B. thailandensis*, *B. glumae*, and *Acinetobacter baumannii* is overactivated upon disruption of *rsaM* homologs, suggesting that putative RsaMs invariably feature as negative regulators of QS circuits (Le Guillouzer et al., 2018; López-Martín et al., 2021; Goo & Hwang, 2023). As *rsaM* mutants in several bacterial species exhibit decreased virulence, this repression has also been linked to maintenance of QS homeostasis (Mattiuzzo et al., 2011; Chen et al., 2012; López-Martín et al., 2021).

**Figure 1.**
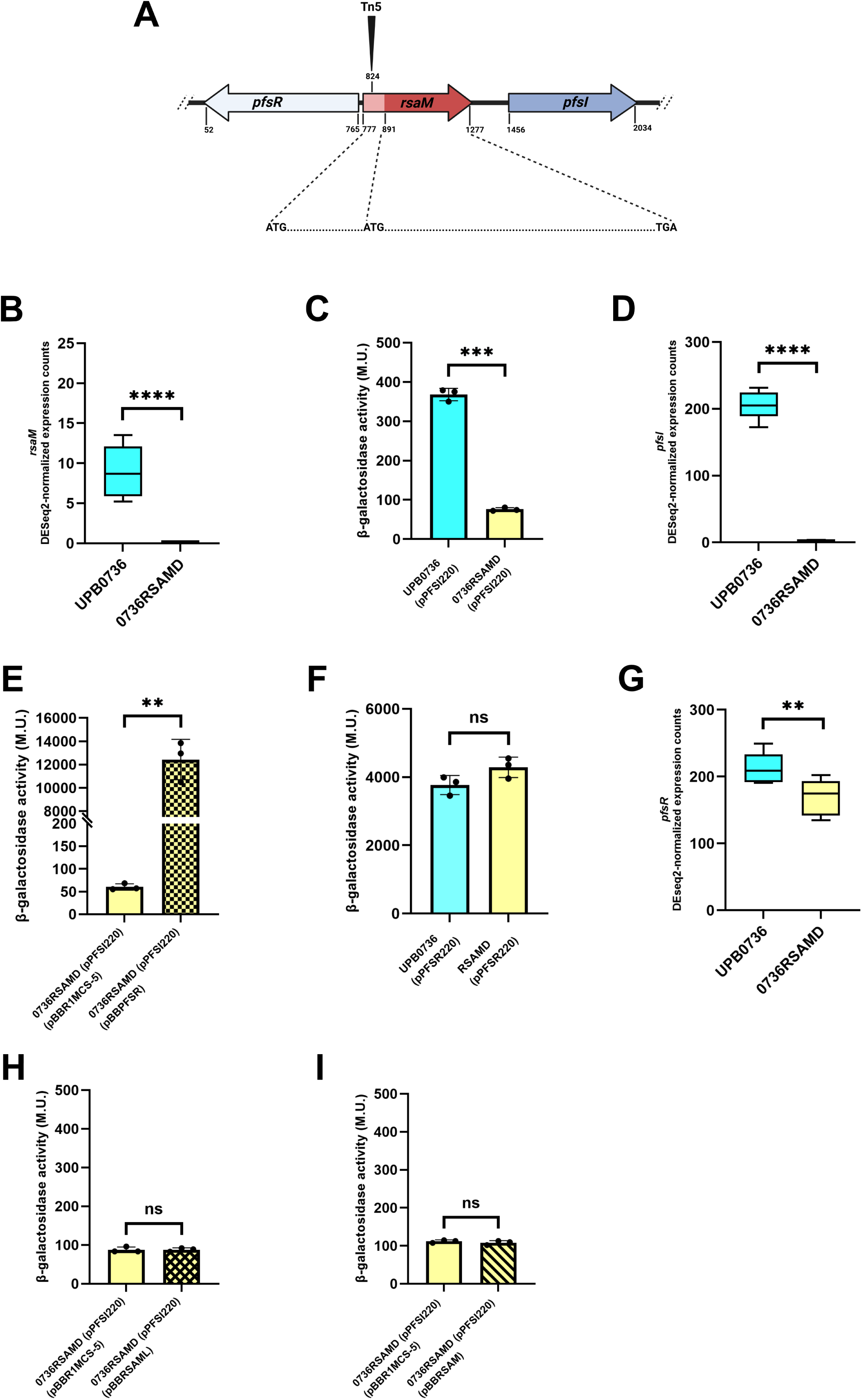
The *rsaM* gene is not required for the maintenance of repression keeping the PfsI/R system silenced. A) Genetic organization of the 2150 bp region coding for *pfsR*, *pfsI* and *rsaM* genes. The shorter ORF of the *rsaM* gene is shown in dark red. Long *rsaM* ORF encompasses the same region as well as the upstream sequence (light red), thus encoding a 38 aa longer protein. The position of the Tn*5* insertion leading to hyperactivation of the PfsI/R system is also indicated. B) DESeq2-normalized counts of *rsaM* transcripts in the wild-type *P. fuscovaginae* UPB0736, obtained through the analysis with the mutant lacking the *rsaM* gene (*P. fuscovaginae* 0736RSAMD). C) *pfsI* promoter activity in *P. fuscovaginae* UPB0736 and 0736RSAMD. D) DESeq2-normalized counts of *pfsI* transcripts in *P. fuscovaginae* UPB0736 and 0736RSAMD. E) *pfsI* promoter activity in *P. fuscovaginae* 0736RSAMD harboring pBBR1MCS-5 or pBBPFSR (*pfsR* under control of *lac* promoter). F) *pfsR* promoter activity in *P. fuscovaginae* UPB0736 and 0736RSAMD. G) DESeq2-normalized counts of *pfsR* transcripts in *P. fuscovaginae* UPB0736 and 0736RSAMD. H) *pfsI* promoter activity in *P. fuscovaginae* 0736RSAMD harboring pBBR1MCS-5 or pBBRSAML (*rsaM* long ORF under control of *lac* promoter). I) *pfsI* promoter activity in *P. fuscovaginae* 0736RSAMD harboring pBBR1MCS-5 or pBBRSAM (*rsaM* shorter ORF under control of *lac* promoter). The promoter activity data are presented as means ± standard deviation. Asterisks indicate significant differences between the means (*p<0.05; **p<0.01; ***p<0.001; ****p<0.0001). All promoter assays were performed with negative controls (test strains carrying empty pMP220); the corresponding β-galactosidase activities are listed in Table S1. The DESeq2­normalized read counts are presented with box plots that depict the medians (central lines), the interquartile ranges (boxes), and minimum and maximum data values for each set of samples (whiskers). Asterisks indicate significant differences between two groups of samples (*p<0.05; **p<0.01; ***p<0.001; ****p<0.0001).

The transposon insertion within the *pfsR/pfsI* intergenic region in *P. fuscovaginae* UPB0736 resulting in decanoyl-homoserine lactone (AHL-C10) hyperproduction was mapped to the possible *N*-terminus-coding region of the longer *rsaM* ORF (Fig. 1A), suggesting that this phenotype could stem from the interruption of the coding sequence of the *rsaM* gene (if long ORF is functional) or *cis*-regulatory elements that are required for its expression (if shorter ORF is functional) (Mattiuzzo et al., 2011). In this study, we confirmed that RsaM is natively expressed, determined which of the two possible *rsaM* ORFs is translated, and then established the functional position of RsaM within the PfsI/R regulatory circuit. RsaM does not serve as a repressor switch of the PfsI/R QS system but instead acts as a negative modulator when this system is triggered by a yet unknown mechanism. Finally, we affirmed that the divergent *pfsR/rsaM* intergenic/promoter region is pivotal for the activation of the PfsI/R system.

## Materials and methods

### Bacterial strains and growth conditions

*E. coli* strains were grown in Luria-Bertani (LB) medium (tryptone 10 g/l, yeast extract 5 g/l, NaCl 10 g/l) supplemented with antibiotics when necessary (ampicillin 100 mg/l, kanamycin 25 mg/l or 100 mg/l, gentamicin 10 mg/l, tetracycline 10 mg/l) at 37 °C with shaking. For overexpression of the recombinant RsaM, *E. coli* M15 carrying the plasmids pQE31RsaM and pREP4 was grown in Terrific broth (TB) medium (tryptone 12 g/l, yeast extract 24 g/l, glycerol 4 g/l, KH_2_PO_4_ 2.3 g/l, K_2_HPO_4_ 12.5 g/l) with added ampicillin 100 mg/l and kanamycin 25 mg/l. *Pseudomonas fuscovaginae* UPB0736 and mutant derivatives were grown in King’s B (KB) medium (proteose peptone no.3 20 g/l, MgSO_4_ x 7H_2_O 1.5 g/l, KH_2_PO_4_ 1.2 g/l, glycerol 10 g/l) at 30° C with shaking. When required, the medium was supplemented with antibiotics (kanamycin 100 mg/l, gentamicin 40 mg/l, tetracycline 40 mg/l, nitrofurantoin 100 mg/l). To make solid media, 13 g/l agar was added.

### Plasmid construction/recombinant DNA techniques

The plasmids and primers used in this study are listed in Table 1 and Table 2, respectively. Routine DNA manipulation techniques, including restriction enzyme digestion, agarose gel electrophoresis, DNA fragment purification, ligation with T4 ligase, PCR, and transformation into *E.coli*, were performed as described previously (Sambrook, 1989).

**Table 1.**
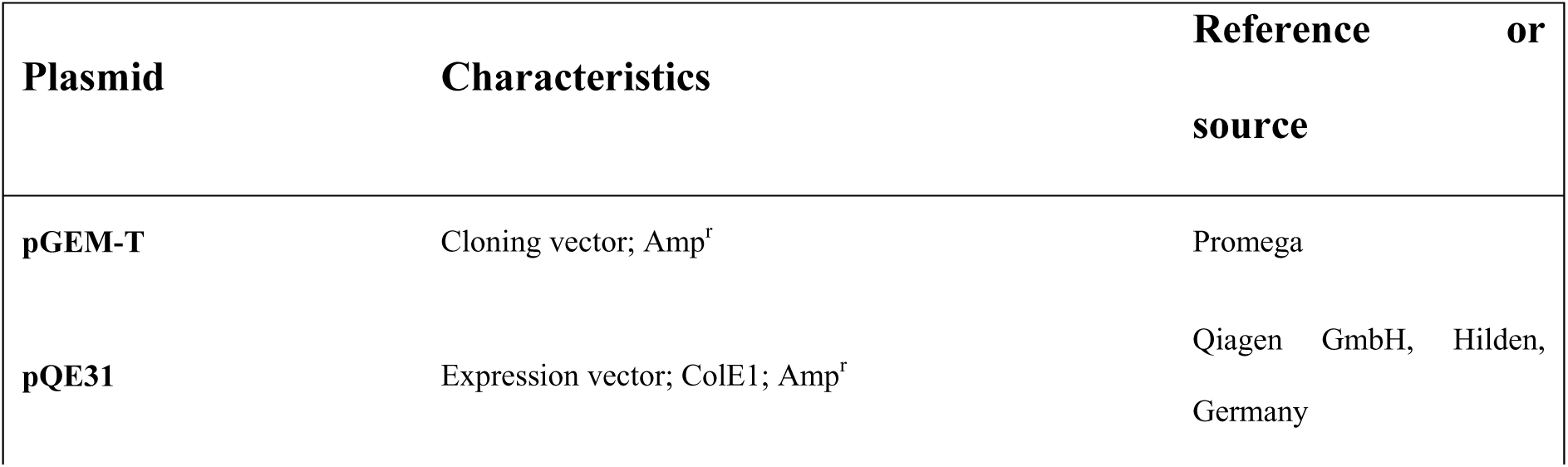
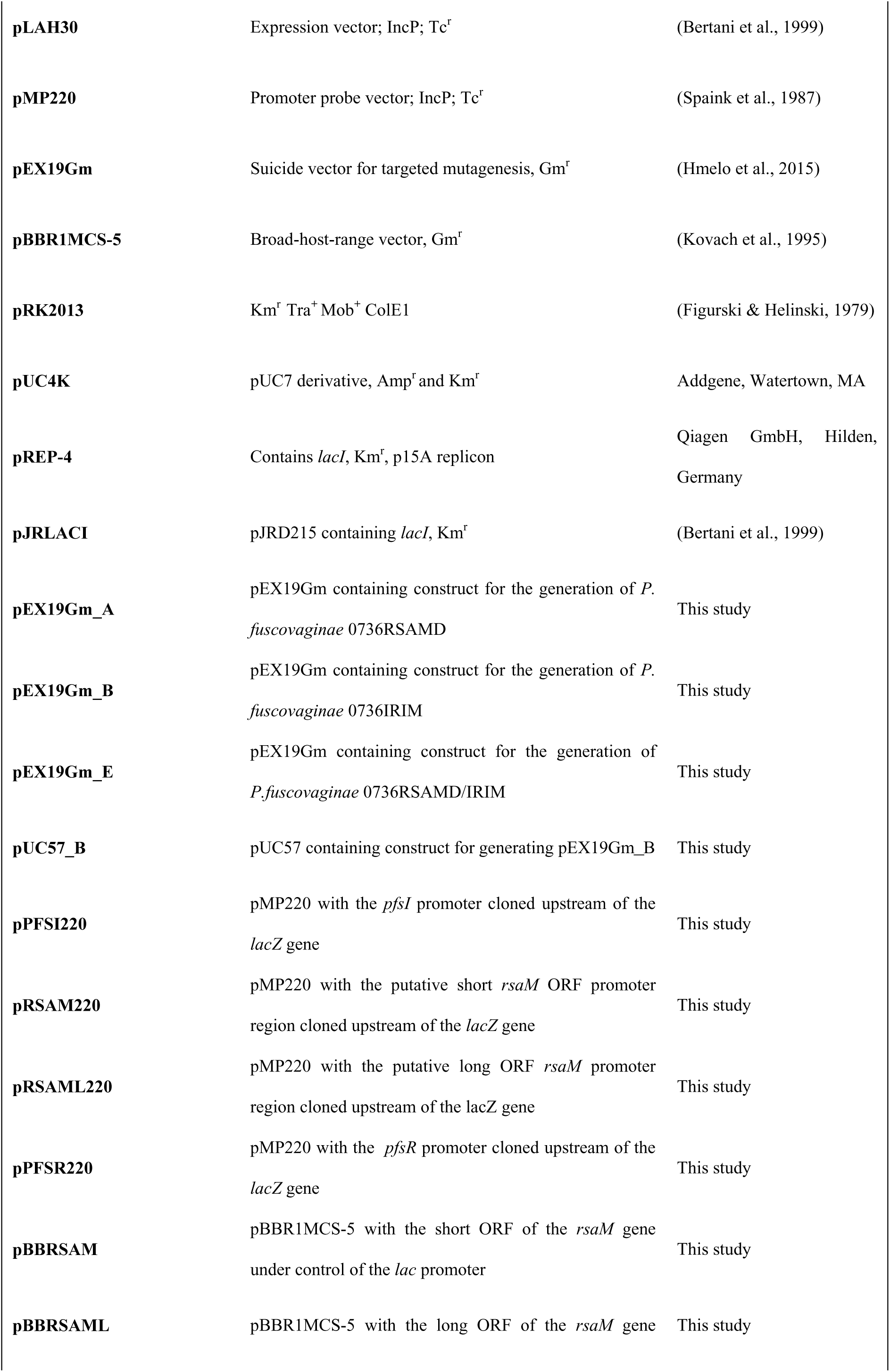
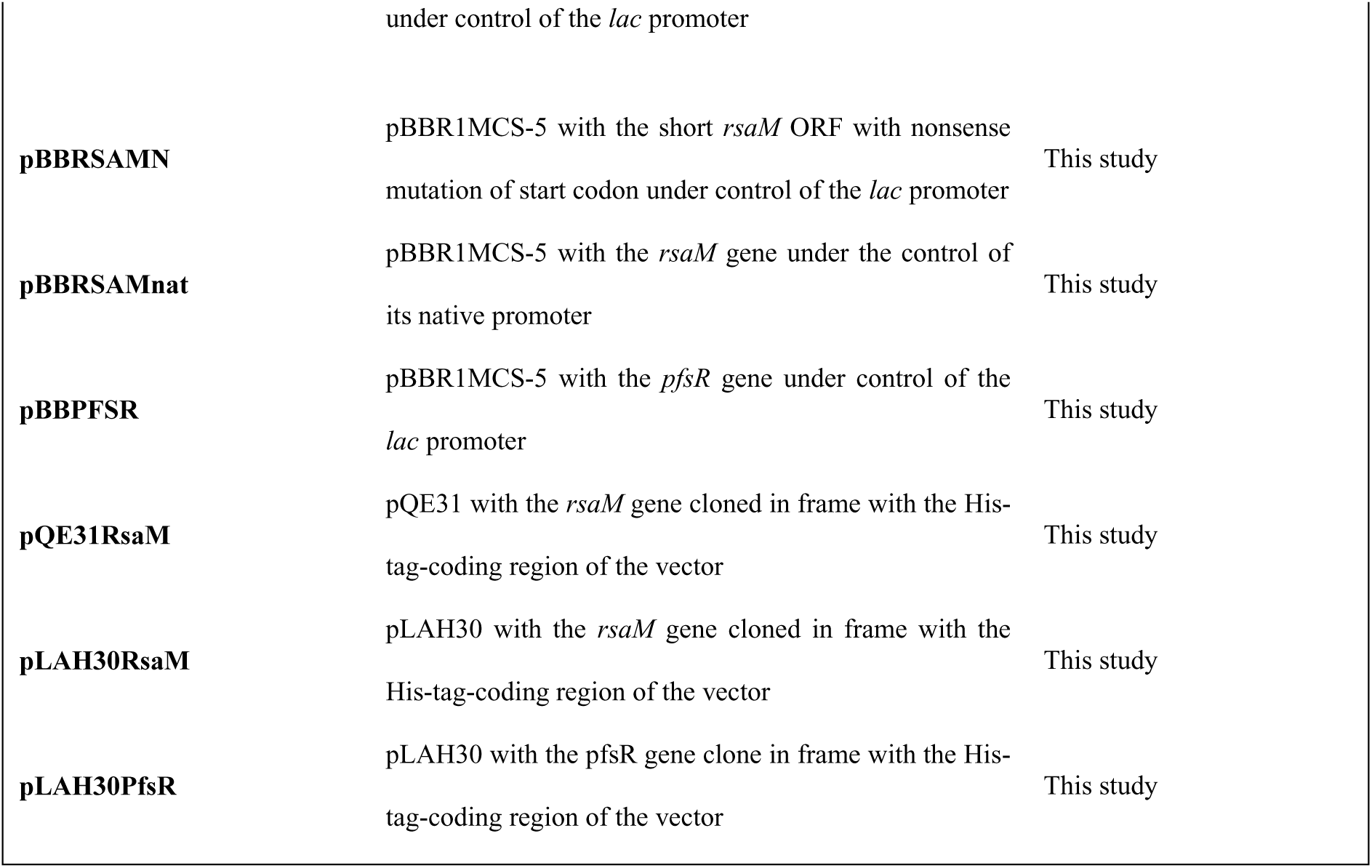
Plasmids used in this study.

**Table 2.**
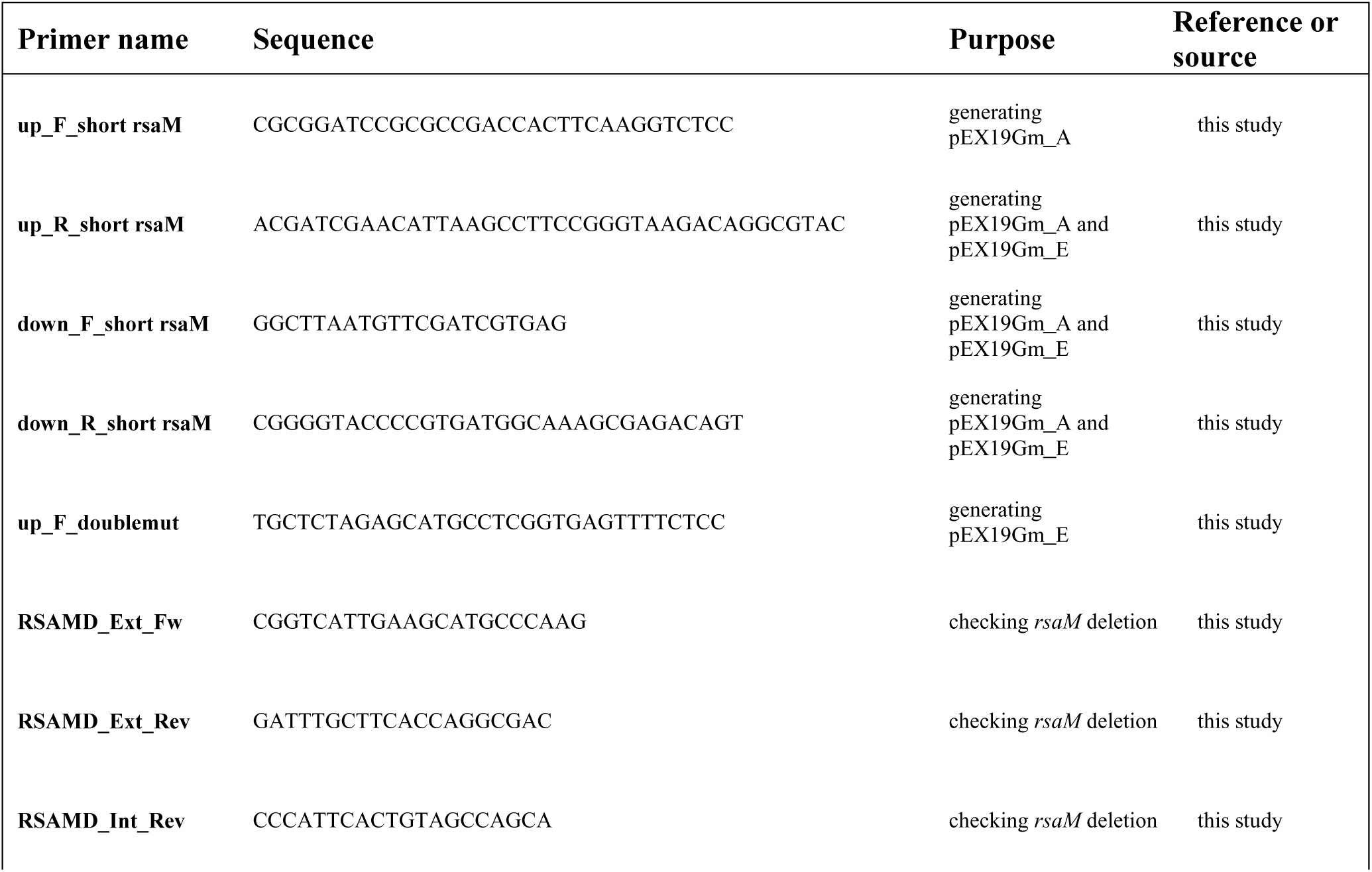
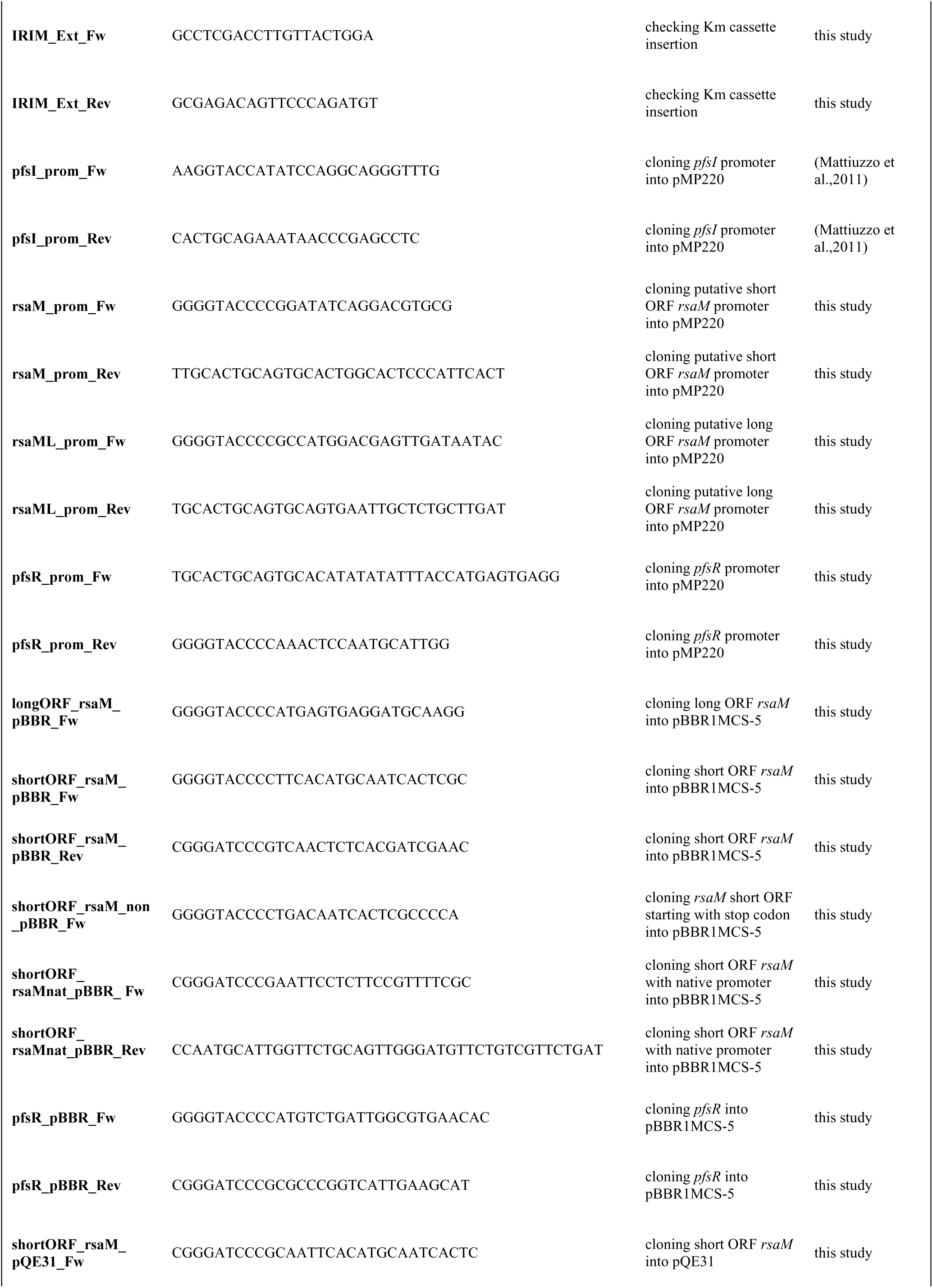
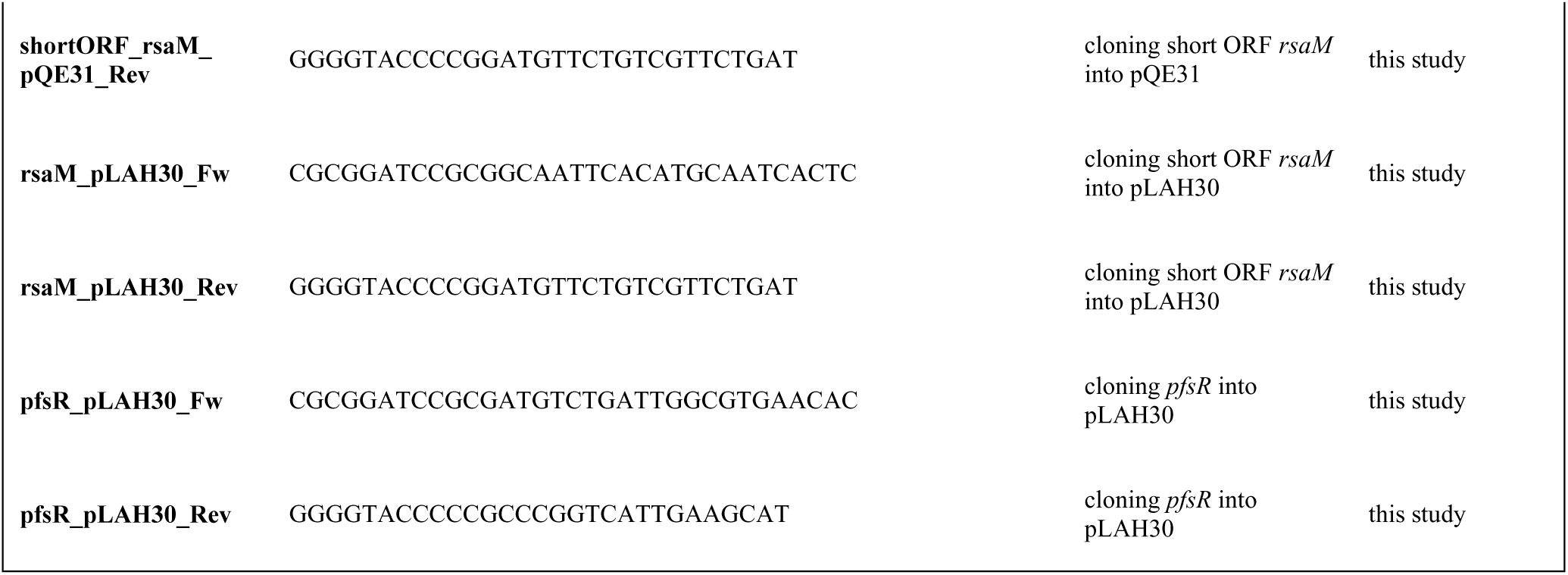
Primers used in this study.

### Generation of *P. fuscovaginae* UPB0736 mutants

Targeted mutagenesis was performed using the suicide vector pEX19Gm, via two-step allelic exchange, as described previously (Hmelo et al., 2015). To generate the *rsaM* deletion mutant in the wild-type background (*P. fuscovaginae* 0736RSAMD), 700 bp upstream and 500 bp downstream of the sequence targeted for removal were first PCR amplified and fused via overlap extension PCR. The fused product was then cloned into pEX19Gm, yielding pEX19Gm_A. To obtain the mutant with the kanamycin cassette inserted in the position of the previously described Tn*5* insertion (*P. fuscovaginae* 0736IRIM), the pUC57 plasmid containing 500 bp upstream and downstream of the target position separated by the BamHI restriction site was synthesized by Genescript. This 1000 bp fragment was cut from the vector and cloned into pEX19Gm. The obtained plasmid was then linearized with BamHI and ligated to the Km cassette excised from the pUC4K plasmid with the same enzyme, thus generating the construct pEX19Gm_B. To obtain the double mutant having both the kanamycin resistance cassette insertion and *rsaM* deletion (*P. fuscovaginae* 0736RSAMD/IRIM), 500 bp upstream and downstream of the target sequence were amplified via PCR using genomic DNA of 0736IRIM as a template. Overlap extension PCR was used to generate the fusion of the two amplicons which was then cloned into pEX19Gm to yield pEX19Gm_E. All constructs were conjugated from *E.coli* DH5α into *P. fuscovaginae* UPB0736 (for mutants 0736RSAMD and 0736IRIM) or *P. fuscovaginae* 0736IRIM (for the mutant 0736RSAMD/IRIM) by triparental mating on solid KB with *E. coli* HB101 harboring the plasmid pRK2013. Conjugation mixtures were incubated overnight at 30°C before transfer onto KB plates containing 50 mg/l nitrofurantoin and 30 mg/l gentamycin to select for transconjugants. The second recombination event was induced by plating putative transconjugants onto NSLB (LB medium without added NaCl) supplemented with 100 mg/l nitrofurantoin and 15% sucrose. Colonies were screened for the presence of deletions or insertions by PCR, using primers with binding sites outside of the region cloned into pEX19Gm. The mutants were confirmed by PCR and sequencing. The genetic maps of the *pfsI/R* locus in *P. fuscovaginae* UPB0736 and mutant strains *P. fuscovaginae* 0736RSAMD, 0736IRIM and 0736RSAMD/IRIM are shown in Table 3.

**Table 3.**
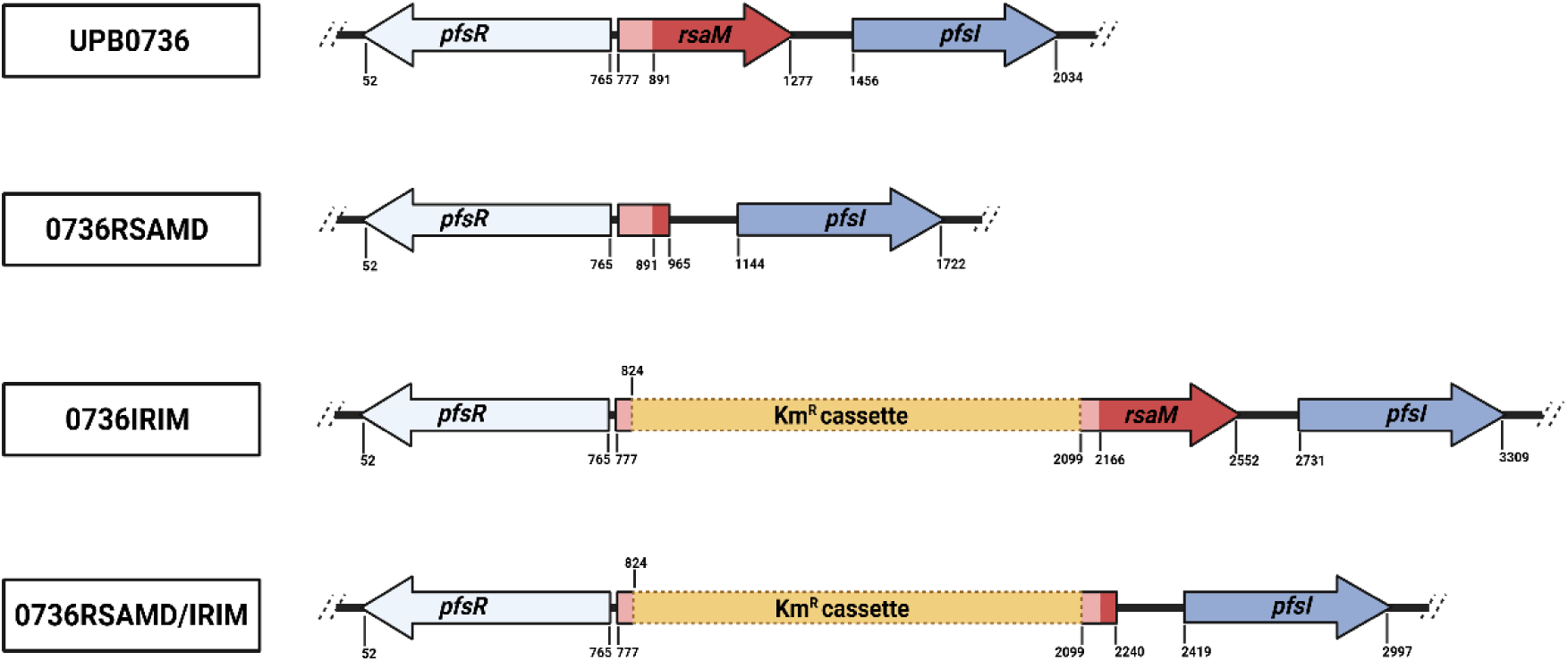
Genetic organization of the *pfsI*/*R* locus in *P. fuscovaginae* UPB0736 (wild-type strain) and mutant derivatives generated in this study: *P. fuscovaginae* 0736RSAMD (*in-frame* deletion of the *rsaM* gene in the wild-type background), *P. fuscovaginae* 0736IRIM (kanamycin resistance cassette inserted at the position of the previously described Tn*5* insertion, and 0736RSAMD/IRIM (*in-frame* deletion of the *rsaM* gene in the mutant harboring kanamycin resistance cassette).

### β-galactosidase promoter activity assays

To assess the activities of the *pfsI*, *pfsR*, and *rsaM* promoters, the corresponding regions were amplified using primer pairs given in Table 2 and genomic DNA of *P. fuscovaginae* UPB0736 as template, followed by cloning into the transcriptional reporter vector pMP220. The constructs were checked for correct sequence and introduced into the wild-type and mutant strains via triparental mating. For the β-galactosidase assay, 5 ml of growth media was inoculated with a single colony of the test strain and incubated overnight for 24 h in flasks. The exception was the test in which the mutant 0736RSAMD harbored constitutively expressed PfsR, which was conducted after 16 h. The tests were conducted essentially as described previously (Miller, 1972), with modifications (Stachel et al., 1985). Negative controls, with test strains carrying empty pMP220, were included in each test. All values shown in the graphs are the averages of three biological replicates, with error bars indicating standard deviation.

### AHLs detection

The production of AHLs was assessed via T-streaks with the AHL biosensor *Chromobacterium violaceum* 026 (McClean et al., 1997). A single colony of each test strain was streaked in line perpendicular to the biosensor strain on KB plates. Plates were incubated for at least 72 h at 30 °C.

### RNA isolation and sequencing

Bacterial cultures were grown in 5 ml of KB medium until the onset of the stationary phase. 1000 µl of RNAprotect Bacteria Reagent (Qiagen) was added to 500 µl of culture (containing approximately 10^9^ CFU), vortexed for 5 s, and centrifuged at 2500xg at 4 °C for 10 min. The supernatant was thoroughly removed, and cell pellets were immediately transferred to dry ice and stored at -80 °C until use. Total RNA was isolated using the RNeasy Mini Kit (Qiagen) following the manufacturer’s instructions with a few modifications. Namely, cells were lysed in TE buffer (Tris 10 mM, EDTA 1 mM, pH 8) with the addition of 5 mg/ml of lysozyme, followed by 20 s of vortexing and 10 min incubation at room temperature. RNA concentration and purity were assessed with Nanodrop, while RNA integrity was verified on a 1.5 % (m/V) agarose gel. RNA sequencing using the Illumina NovaSeq X Plus Sequencing System, as well as the subsequent bioinformatic and statistical analysis, was conducted by Novogene. The sequencing data were deposited in NCBI Bioproject under the accession number PRJNA1370452.

### Recombinant protein expression, purification and anti-RsaM antibodies generation

An overnight culture of *E. coli* M15 harboring the plasmids pREP4 and pQE31RsaM was diluted in 250 ml fresh TB medium (starting OD_600_ 0.05). Upon reaching the mid-exponential phase (OD_600_ 0.6), IPTG was added to the culture (final concentration 1 mM) which was then incubated for 5 h (37 °C, with shaking). The culture was centrifuged at 2500xg at 4 °C for 25 min, and the pellets were stored at 4 °C until purification. Since the recombinant RsaM localized entirely in the insoluble fraction of the lysate, a protocol for the purification of inclusion bodies was used prior to affinity purification. Briefly, the cells were resuspended in 1 mM dithiothreitol (DTT) and disrupted using a French press (2.5 kbar, 4 °C), followed by centrifugation of the lysate at 9300xg at 4 °C for 30 min. The pellet was washed two times in a buffer (50 mM Tris-HCl buffer, 5 mM DTT, 5 mM EDTA, 0.75 % (m/V) sodium deoxycholate, pH 8) and once in water, each time followed by centrifugation at 9300xg at 4 °C for 30 min. The purified inclusion body was solubilized in Tris-urea buffer (10 mM Tris, 8 mM urea, 100 mM NaH_2_PO_4,_ pH 8), and the recombinant RsaM was purified under denaturing conditions using the QIAexpressionist kit (Qiagen), according to the manufacturer’s instructions. Bradford reagent (Biorad) was used to determine protein concentration. The expression of the recombinant protein and its presence throughout purification were confirmed via standard SDS-PAGE, as described previously (Laemmli, 1970). Anti-RsaM antibodies were raised in rabbits and affinity-purified at Thermofisher.

### Western blot and immunoprecipitation

To obtain samples for Western blots, 1 ml of each culture (grown 4 h, 16 h or 24 h) was centrifuged at 3300xg for 5 min, followed by resuspension of the cell pellets in 10 cell volumes of sample buffer (4 % SDS, 20 mM DTT, 0.01% bromophenol blue, 60 mM Tris, pH 8), boiling (95 °C, 5 min), sonication (30 sec at full force), and centrifugation at 9300xg for 10 min. Equal volumes of cleared lysate (7.5 µl) were loaded onto a discontinuous SDS PAGE gel (5% stacking, 12% resolving). Separated proteins were transferred onto a PVDF membrane activated with PBST buffer with added methanol (1xPBS, 0.1% Tween, 10% methanol) via wet transfer. The membrane was then incubated in a PBST buffer containing 5% skimmed milk (2 h at room temperature), followed by incubation with anti-RsaM antibodies (1:500 dilution in PBST) overnight at 4 °C. The membrane was then washed three times with PBST buffer (7 min each wash), and incubated with the 1:2500 dilution of the anti-IgG-rabbit HRP-conjugated secondary antibody for 1 h at room temperature. After three washes with PBST, the membrane was incubated with ECL (BioRad), as described by the manufacturer. Pictures of the membrane were taken on ChemiDoc (BioRad). For immunoprecipitation, 1 ml of the overnight cultures (24 h) was centrifuged (2500xg, 4 °C, 5 min). The pellets were washed once in PBS, resuspended in 1 ml of the RIPA buffer (150 mM NaCl, 1% NP-40, 0.5% sodium deoxycholate, 0.1% SDS, 50 mM Tris, pH 8), followed by sonication (on ice, 20 s at maximum intensity, four times with 1 min pauses). The lysate was cleared by centrifugation at 10000xg, at 4 °C, for 10 min, passed through a 0.22 µM filter, and incubated with the affinity-purified anti-RsaM antibodies (final concentration 500 µg/ml) for 1 h at 4 °C. The EZview^TM^ Protein G Affinity Gel (Sigma) was then added to the lysate, with the rest of the immunoprecipitation protocol performed according to the manufacturer’s instructions.

### Statistical analysis

Welch’s T-tests were used to compare the means obtained in promoter assays. The analysis was performed in Prism 10.4.2 (GraphPad Software).

## Results

### The *rsaM* gene is not expressed in the wild-type *P. fuscovaginae*, and it is not required for maintaining the PfsI/R system silent

We hypothesized that overproduction of PfsI-synthesized AHLs in the previously reported *rsaM*::Tn*5* mutant (Mattiuzzo et al., 2011) was either due to the interruption of the *rsaM* coding sequence or of the *rsaM-pfsR* divergent intergenic/promoter region. The *rsaM* gene has two possible ORFs in the same frame, one having a 38 amino acid extension at the *N*-terminus (Fig. 1A). It was therefore initially of interest to confirm the presence of RsaM in the wild-type *P. fuscovaginae* and elucidate the correct ORF. To do this, we cloned the shorter ORF of *rsaM* into an expression vector and successfully expressed the His-tagged protein in *E. coli*. The recombinant protein was purified and used to raise polyclonal antibodies as described in the Materials and Methods section. The antibodies were then used in Western blot analysis and immunoprecipitation coupled with mass spectrometry (IP-MS). Surprisingly, RsaM was not detected in the wild-type *P. fuscovaginae* with either technique (data not shown). In line with this, putative promoters of the *rsaM* gene showed low activity (putative promoter for the short *rsaM* ORF) or no activity (putative promoter for the long *rsaM* ORF) (Fig. S1) while the transcriptomic analysis evidenced an extremely low level of the *rsaM* transcription (including transcripts originating from the extension of the long *rsaM* ORF) (Fig. 1B). It was concluded that *rsaM* is not transcribed in the wild-type strain and no RsaM protein is present in the cell.

To determine whether loss of function of the *rsaM* gene causes PfsI/R overactivation, targeted mutagenesis was performed by generating an *in-frame* deletion mutant where most of the shorter *rsaM* ORF (and thus also the long ORF) was deleted (Table 3). The resulting mutant, *P. fuscovaginae* 0736RSAMD, did not produce AHLs, displayed no *pfsI* promoter activity, and lacked *pfsI* transcripts. This was in contrast with the wild type where very low levels of AHL production and *pfsI* transcription were present (Fig. S2; Fig. 1C and D). To confirm that the deletion of the *rsaM* gene caused no *cis*-acting changes to the probable *lux* box upstream of *pfsI* (and downstream of the *rsaM* ORFs), we constitutively expressed PfsR *in trans* in this mutant. This resulted in the activation of the *pfsI* promoter, evidencing that the *lux* box sequence, and hence *pfsI* transcriptional expression, remained functional (Fig. 1E). The *pfsR* transcript level decreased only slightly in this mutant compared to the wild type; therefore, the cessation of *pfsI* transcription most probably was not due to changes in the level of PfsR (Fig. 1F and G). We were not able to complement the phenotype of this mutant upon introduction of the long or short *rsaM* ORF under control of a constitutively active *lac* promoter *in trans*; the reason for this is currently unknown (Fig. 1H and I). Taken together, these results indicated that the *rsaM* gene is not transcribed in the wild-type *P. fuscovaginae* under the laboratory conditions used and that this locus does not exert a top-level repressive effect on the PfsI/R QS system.

### The intergenic region between *pfsR* and *rsaM* is key in triggering the PfsI/R system

As we established that the repression of the PfsI/R system is not the direct effect of the *rsaM* gene, we hypothesized that the *rsaM*::Tn*5* insertion (Mattiuzzo et al., 2011) caused hyperactivation of the circuit by disrupting another regulatory element. To further validate this effect/phenotype in the *rsaM*::Tn*5* mutant (Fig. 1A), we regenerated this mutant by introducing a kanamycin resistance cassette in the exact genomic position of the Tn*5* integration (Table 3). The resulting mutant, *P. fuscovaginae* 0736IRIM, displayed a similar phenotype as the *rsaM*::Tn*5* mutant, having considerably elevated AHL production as well as significantly elevated *pfsR* and *pfsI* promoter activities (and corresponding RNA transcripts as determined from the RNAseq data) with respect to the wild type *P. fuscovaginae* (Fig. S3; Fig. 2A, B, C and D). Remarkably, the transcription of the *rsaM* gene in this mutant was also significantly increased compared to the wild type, as evidenced by promoter studies and transcriptomic analysis (Fig. 2E and F). Consistent with this, both Western blot analysis and IP-MS showed that the mutant expressed the RsaM protein corresponding to the predicted shorter ORF-encoded form (Fig. 2G and H, Fig. S4). To our knowledge, this is the first evidence of a protein belonging to the RsaM family being natively expressed. These results led us to conclude that the shorter ORF of the *rsaM* gene is the functional ORF and that insertions in the regulatory *pfsR-rsaM* divergent promoter region activate the transcription of both *pfsR* and *rsaM* genes.

**Figure 2.**
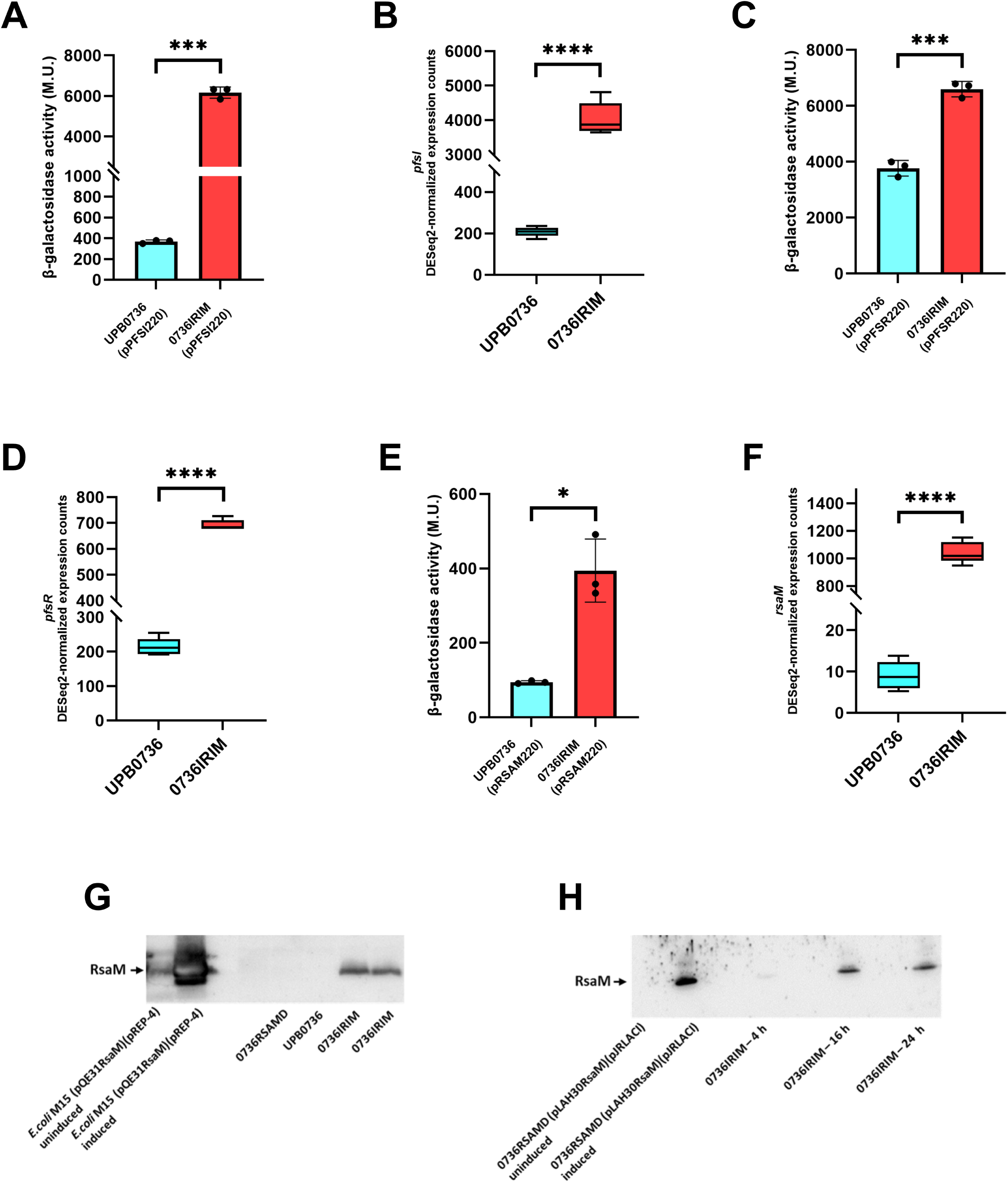
The intergenic *pfsR*-*rsaM* region is crucial in repressing the *pfsI*, *pfsR*, and *rsaM* genes. A) *pfsI* promoter activity in *P. fuscovaginae* UPB0736 and 0736IRIM. B) DESeq2-normalized counts of *pfsI* transcripts in *P. fuscovaginae* UPB0736 and 0736IRIM. C) *pfsR* promoter activity in the *P. fuscovaginae* UPB0736 and 0736IRIM. D) DESeq2-normalized counts of *pfsR* transcripts in *P. fuscovaginae* UPB0736 and 0736IRIM. E) *rsaM* promoter activity (short *rsaM* ORF promoter) in the *P. fuscovaginae* UPB0736 and 0736IRIM. F) DESeq2-normalized counts of *rsaM* transcripts in *P. fuscovaginae* UPB0736 and 0736IRIM. G) Western blots performed with anti-RsaM antibodies on the total lysates of *E. coli* M15 (pREP4) harboring pQE31RsaM without (lane 1) or with induction with IPTG (lane 2), *P. fuscovaginae* 0736RSAMD (lane 5), wild-type *P. fuscovaginae* UPB0736 (lane 6) and *P. fuscovaginae* 0736IRIM (lanes 7 and 8). H) Western blots performed with anti-RsaM antibodies on the total lysates of *P. fuscovaginae* 0736RSAMD (pJRLACI) harboring pLAH30RsaM without (lane 1) or with added IPTG (lane 2), *P. fuscovaginae* 0736IRIM in late exponential phase (lane 5), mid-stationary phase (lane 8), and late stationary phase (lane 11). Lysates of the wild-type strain and *P. fuscovaginae* 0736RSAMD obtained from the same timepoints are in the lanes 3, 6, 9 and 4, 7, 10, respectively. Full pictures of Western blots are shown in the Supplementary (Fig. S13). The promoter activity data are presented as means ± standard deviation. Asterisks indicate significant differences between the means (*p<0.05; **p<0.01; ***p<0.001; ****p<0.0001). All promoter assays were performed with negative controls (test strains carrying empty pMP220); the corresponding β-galactosidase activities are listed in Table S1. The DESeq2­normalized read counts are presented with box plots that depict the medians (central lines), the interquartile ranges (boxes), and minimum and maximum data values for each set of samples (whiskers). Asterisks indicate significant differences between two groups of samples (*p<0.05; **p<0.01; ***p<0.001; ****p<0.0001).

### RsaM is a post-activation negative modulator, not the master repressor, of the PfsI/R system

To gain insight into the regulatory role of RsaM, we first introduced short ORF *rsaM* cloned into a plasmid into the insertional mutant 0736IRIM, either under the control of the native *rsaM* promoter or the *lac* promoter present on the backbone of the vector pBBR1MCS-5. Introduction of the constitutively expressed *rsaM* gene resulted in approximately a twofold decrease in *pfsI* promoter activity, and a significant decrease in AHL production (Fig. S5; Fig. 3A). Albeit not statistically significant, we also observed a decrease in *pfsI* promoter activity when the *rsaM* gene was expressed under its own promoter (Fig. 3B). Based on these results, we hypothesized that RsaM constrains activation of the PfsI/R system when triggered by the disruption of the *pfsR*-*rsaM* intergenic region. To further examine this, we carried out targeted mutagenesis and deleted the *rsaM* gene in the 0736IRIM mutant, thereby attaining activation of the PfsI/R system in the absence of RsaM (Table 3). The obtained double mutant (*P. fuscovaginae* 0736RSAMD/IRIM) produced significantly higher levels of AHLs compared to the mutant 0736IRIM (Fig. S6). This effect could be complemented since the production of AHLs decreased dramatically upon introduction of the shorter ORF *rsaM* under control of the *lac* promoter *in trans* (Fig. S7), as well as *pfsI* promoter activity (Fig. 3C). To verify that the complementation observed here was directly caused by the RsaM protein, we also generated a construct with *lac* promoter-controlled shorter *rsaM* ORF having a nonsense mutation of the start codon and introduced it into this mutant. AHL production was not affected in the presence of this construct (Fig. S8), indicating that trans effects of the intact shorter *rsaM* ORF under the control of the same promoter are exclusively RsaM protein-mediated. Overall, these results evidenced that RsaM functions as an intrinsic negative modulator that limits the activation of the PfsI/R circuit once the system has been triggered. Interestingly, despite the large difference in AHL production, there was no significant change in *pfsI* promoter activity between the double mutant and its parent mutant strain (Fig. 3D), suggesting that RsaM possibly functions posttranscriptionally.

**Figure 3.**
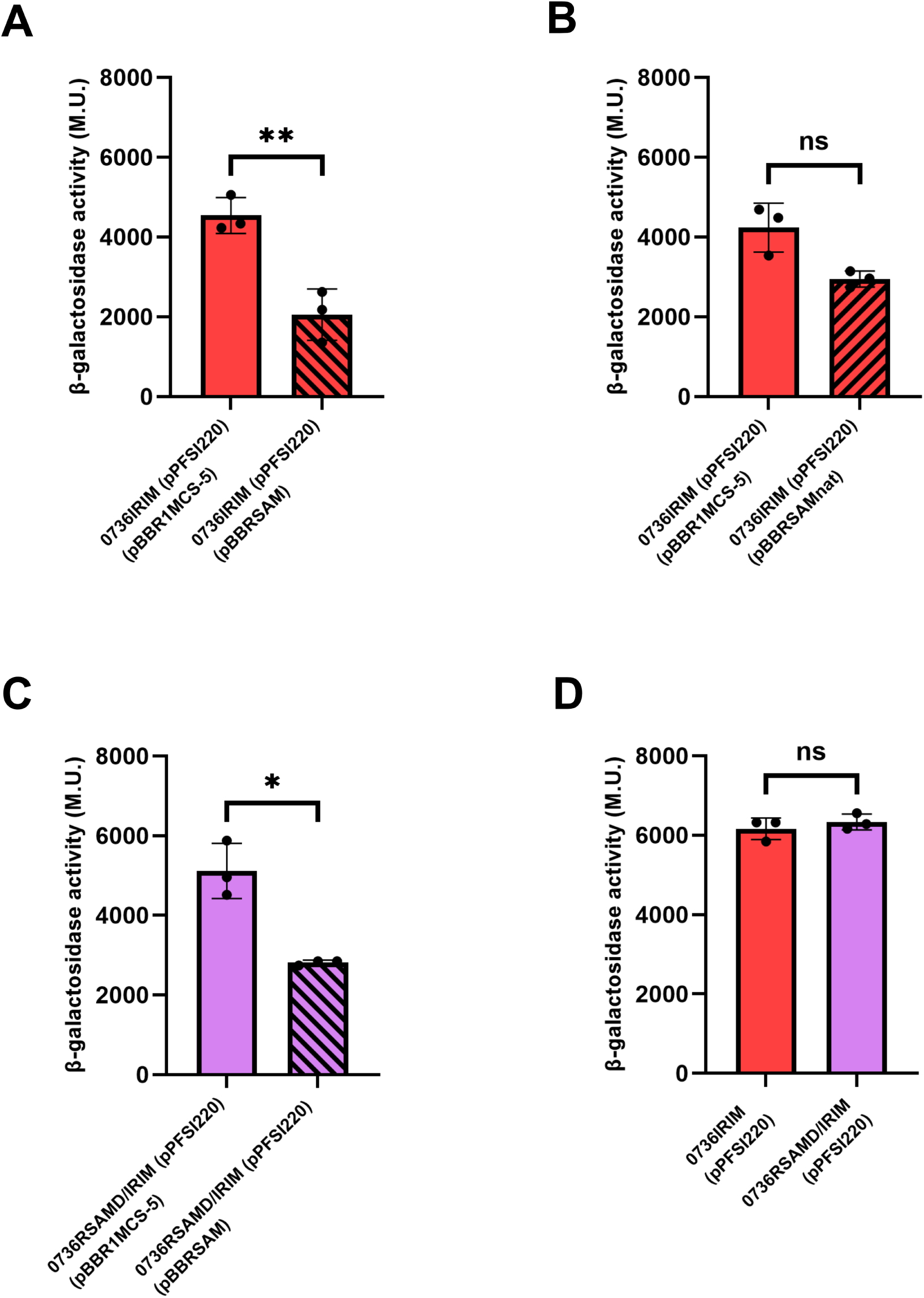
RsaM is a negative modulator of the activated PfsI/R system. A) *pfsI* promoter activity in *P. fuscovaginae* 0736IRIM harbouring pBBR1MCS-5 (control) or pBBRSAM (short ORF *rsaM* under control of *lac* promoter). B) *pfsI* promoter activity in *P. fuscovaginae* 0736IRIM with pBBR1MCS-5 (control) or pBBRSAMnat (short ORF *rsaM* under control of the native promoter). C) *pfsI* promoter activity in *P. fuscovaginae* 0736RSAMD/IRIM with pBBR1MCS-5 (control) or pBBRSAM (short ORF *rsaM* under control of *lac* promoter). D) *pfsI* promoter activity in *P. fuscovaginae* 0736IRIM and 0736RSAMD/IRIM. The promoter activity data are presented as means ± standard deviation. Asterisks indicate significant differences between the means (*p<0.05; **p<0.01; ***p<0.001; ****p<0.0001). All promoter assays were performed with negative controls (test strains carrying empty pMP220); the corresponding β-galactosidase activities are listed in Table S1.

Following these findings, it was of interest to investigate whether RsaM is expressed and modulates the activity of the PfsI/R system in the wild-type *P. fuscovaginae* when this circuit is triggered by overexpressing PfsR *in trans*. RsaM was not detected in the wild type via Western blots under these conditions, suggesting that PfsR alone is not sufficient to activate the transcription of the *rsaM* gene (not shown). These results were surprising given that the wild-type strain harbouring the *pfsR* gene *in trans* under a constitutive promoter produced a lower amount of AHLs than the *rsaM* mutant, 0736RSAMD, carrying the same construct (Fig. S9). Since endogenous RsaM was expressed only upon interruption of the *pfsR-rsaM* intergenic region, it is possible that the same repression mechanism silencing the *pfsR* gene also acts on the expression of the *rsaM* gene; whether this is the case remains to be established. With this in mind, we also asked if the *rsaM* gene expressed from the *lac* promoter *in trans* in the wild-type *P. fuscovaginae* UPB0736 would alter PfsI/R QS dynamics when the strain is constitutively expressing PfsR, thus achieving simulation of the 0736IRIM mutant background. RsaM-mediated inhibition/modulation of the circuit was once again evident, consolidating the results of the studies reported above (Fig. S10).

## Discussion

*P. fuscovaginae* possesses two archetypical QS systems that are silenced *in vitro* under standard growth conditions, but are activated *in planta*, where they play a role in virulence. A Tn*5* insertion in the *pfsI*-*pfsR* intergenic region was previously found to hyperactivate the PfsI/R system, raising the hypothesis that the uncharacterized regulatory protein RsaM, encoded in this region, might play a repressive role in regulating this circuit (Mattiuzzo et al., 2011). The results we obtained here revealed that RsaM is absent in the wild-type *P. fuscovaginae* and thus does not contribute to blocking the activation of the PfsI/R system in laboratory settings. We demonstrated that an insertion in the position of Tn*5* integration that triggers the PfsI/R system does not disrupt the functional ORF of the *rsaM* gene but instead activates its expression, and that RsaM acts as a built-in negative modulator of this circuit.

RsaM was not detected in the wild-type *P. fuscovaginae* under culture conditions used in this study, either via Western blot analysis or IP-MS, and the transcription level of the *rsaM* gene was found to be extremely low. As many bacterial regulators are known to function in a low number of copies per cell, it could not be excluded that this is the case with RsaM as well, with the protein amount being below the detection limit of the techniques used in this study (McAdams & Arkin, 1999; Pulkkinen & Metzler, 2015). However, an *in-frame* deletion of *rsaM* in the wild-type background did not result in an AHL-overproducing phenotype, suggesting this gene is not pivotal in silencing the PfsI/R circuit. Due to the loss of *pfsI* expression, the mutant did not produce any detectable AHLs. This phenotype and the lack of complementation upon introduction of the intact *rsaM* gene *in trans* raised the possibility that the deletion interrupted a *cis*-regulatory element controlling the *pfsI* transcription. Interestingly, an *in-frame* deletion of the *tofM* gene, the *rsaM* homologue in *Burkholderia glumae* 336gr-1, resulted in a small increase in AHL production that could not be complemented by expressing TofM *in trans* (Chen et al., 2012).

The results of this study affirmed that the *pfsR*-*rsaM* intergenic divergent promoter region is critical in maintaining the PfsI/R system in a quiescent state. Activation of the system in the insertional mutant in the intergenic region is presumably triggered by PfsR, which would likely accumulate as a result of increased transcription of the *pfsR* gene. Contrary to our initial hypothesis, the disruption of this intergenic region did not inactivate the *rsaM* gene, but instead triggered its expression. Deletion of the *rsaM* gene in this genetic background conferred a markedly different phenotype compared to the same deletion in the wild-type strain, as this mutant hyperproduced AHLs, a phenotype that was complemented (i.e, reducing AHL production) upon constitutively expressing RsaM *in trans*. It can therefore be concluded that once the PfsI/R is activated, RsaM primarily functions as a negative modulator within the PfsI/R system that adjusts the output of this circuit rather than as a top-level repressor blocking its activation. The mode of action of RsaM is currently unknown; however, it is possible that it takes place post-transcriptionally since the *rsaM* deletion in the mutant 0736IRIM did not result in a change in *pfsI* promoter activity. It remains to be determined if RsaM functions similarly to TofM in *Burkholderia glumae* BGR1, which was recently reported to potentially bind the 5’UTR of the synthase *tofI* mRNA and block its translation. Unlike TofM, RsaM does not display significant homology to the RNA-binding protein RsmA (Goo & Hwang, 2023).

The regulatory role of RsaM appears to be similar to that of RsaL, the built-in DNA-binding repressor of the LasI/R system in *P. aeruginosa* and of the PpuI/R system of *P. capeferrum* that counteracts LuxR-family driven transcriptional activation of the synthase gene, thereby restricting the magnitude of the QS response (Rampioni et al., 2007). RsaL is found across *Pseudomonas* and *Burkholderia* species having LasI/R-like systems and is typically encoded by a gene embedded between the *lasI*-*lasR* gene pair. RsaL inactivation results in considerable overproduction of AHLs (Bertani & Venturi, 2004; Rampioni et al., 2007; Suárez-Moreno et al., 2008; Venturi et al., 2011).

Our results evidenced that the *pfsR*-*rsaM* intergenic region plays a major role in repressing the PfsI/R system in the wild-type *P. fuscovaginae*, independently of RsaM. It can be hypothesized that this region contains a binding site for a repressor that functions as a master regulator responsible for superimposing repression on the PfsI/R system and that such a regulator acts bidirectionally, repressing the transcription of both the *pfsR* and *rsaM* genes. The large insertion introduced in this study could have impaired the function of this protein by distorting its binding site or by creating a spacer that impacted the position of the regulator relative to the target promoters (Huo et al., 2009; Levo et al., 2015; Sangeeta et al., 2024). Studies on *rsaM* homologues in *A. baumannii* AB5705 and *B. glumae* BGR1 reported that transposon insertions within the *abaM* and *tofM* genes result in AHL hyperproduction (López-Martín et al., 2021; Goo & Hwang, 2023). To address this, we also constructed a mutant having a kanamycin resistance cassette inserted after the third codon of the short *rsaM* ORF (Fig. S11), which phenocopied the double mutant, indicating elimination of both the putative master regulator- and RsaM-mediated repression. A frameshift mutation at the same position did not cause a drastic change in AHL production (Fig. S12), reaffirming that RsaM alone is not involved in imposing silencing on the PfsI/R system, and suggesting this small disruption is not sufficient to perturb the function of the unknown repressor.

QS can impose a substantial fitness burden on quorate populations, depending on the types and number of the target loci it regulates (Pai et al., 2012; Ruparell et al., 2016). The removal of repressive elements of the QS circuits often deregulates loci responsible for phenotypes that are timely and beneficial for bacterial populations in their ecological niches (Morici et al., 2007; Venturi et al., 2011; Gupta & Schuster, 2013). It is possible that a high fitness cost of QS activation could have driven the evolution of a regulatory organization involving multi-layered repression and modulation in *P. fuscovaginae*. Given their role in virulence, it is probable that QS systems in *P. fuscovaginae* are triggered when the bacterial population encounters specific conditions *in planta*. The signals required for alleviating the superimposed repression and regulators transducing these signals to the PfsI/R circuit, as well as their mechanism of action, remain to be investigated. Once this level of repression is hampered, RsaM likely acts as a control mechanism countering the PfsR-driven positive feedback loop that ensures the magnitude of QS activation does not cross the levels that would significantly compromise fitness (Spacapan et al., 2023).

A model of the regulatory architecture of the PfsI/R system in *P. fuscovaginae* based on the results obtained in this study is depicted in Figure 4. A hypothetical master regulator of the PfsI/R system likely carries out the stringent repressive function that was previously attributed to RsaM, silencing the transcription of all three genes within the *pfsI*/*pfsR* locus. The system is activated most likely in response to cell numbers/density and an unidentified signal/s, with the hypothetical repressor possibly functioning as the switch that unlocks the PfsI/R system. Once the repression is alleviated, the expression of RsaM is also activated. In turn, RsaM acts on one of the elements of the circuit, modulating the activity of the system.

**Figure 4.**
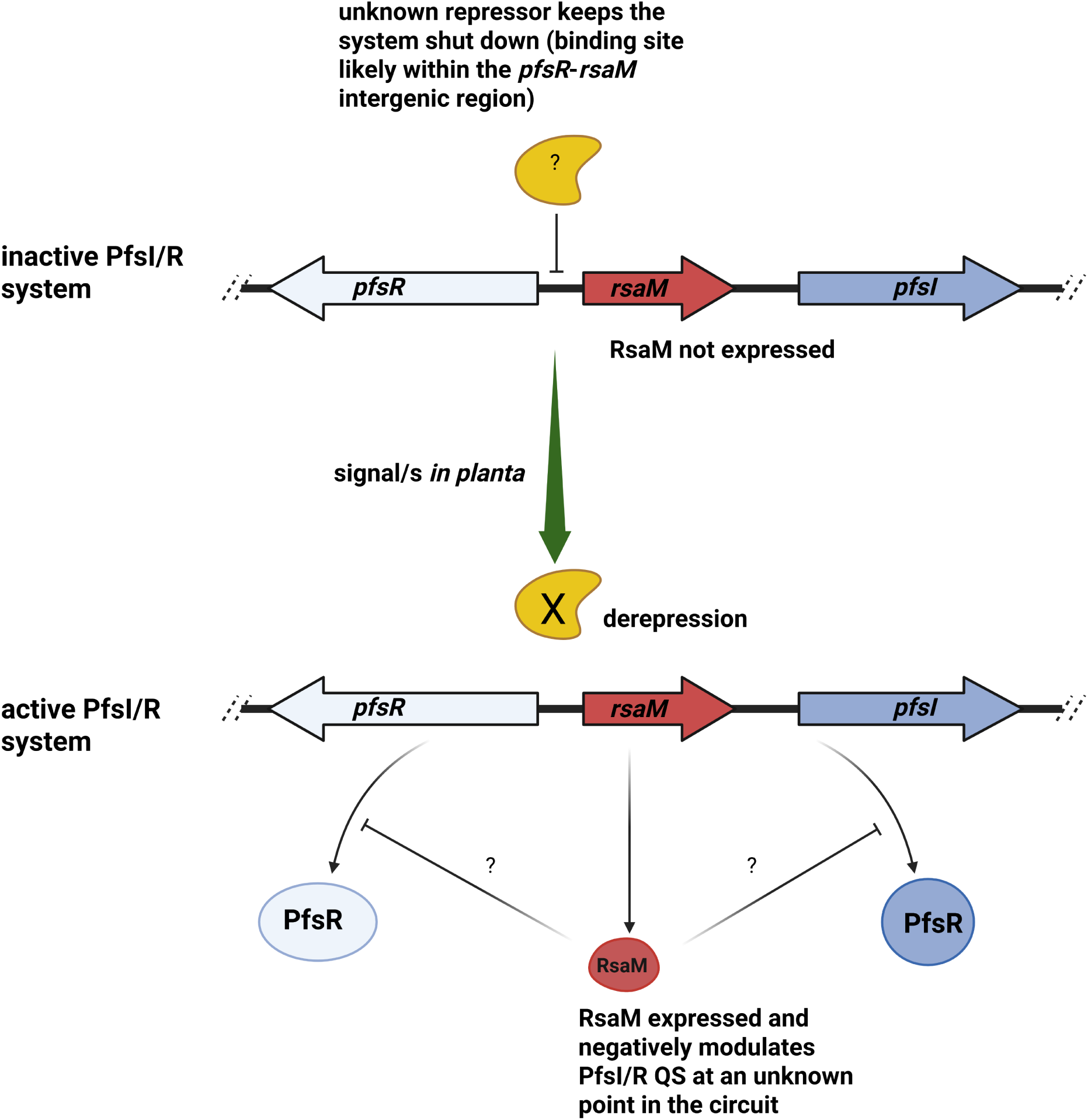
Current model of the regulatory architecture of the PfsI/R system.

## Acknowledgements

N.R. is a recipient of an ICGEB Arturo Falaschi Fellowship, and C.B. from the European Union’s Horizon Europe research and innovation programme under the Marie Skłodowska-Curie grant agreement No 101150379.

## Authors contributions

N.R. (Conceptualization, Data curation, Formal Analysis, Investigation, Methodology, Visualization, Writing – original draft, Writing –review & editing), I.B. (Project administration, Supervision), G.T. (Methodology, Validation), M.P.M. (Methodology, Data curation, Validation), C.B. (Conceptualization, Data curation, Investigation, Supervision, Validation, Writing – original draft, Writing – review & editing), V.V. (Conceptualization, Funding acquisition, Investigation, Project administration, Resources, Supervision, Validation, Writing – original draft, Writing – review & editing).

